# Study of Oak Ridge soils using BONCAT-FACS-Seq reveals that a large fraction of the soil microbiome is active

**DOI:** 10.1101/404087

**Authors:** Estelle Couradeau, Joelle Sasse, Danielle Goudeau, Nandita Nath, Terry C. Hazen, Benjamin P. Bowen, Rex R. Malmstrom, Trent R. Northen

## Abstract

The ability to link soil microbial diversity to soil processes requires technologies that differentiate active subpopulations of microbes from so-called relic DNA and dormant cells. Measures of microbial activity based on various techniques including DNA labelling have suggested that most cells in soils are inactive, a fact that has been difficult to reconcile with observed high levels of bulk soil activities. We hypothesized that measures of *in situ* DNA synthesis may be missing the soil microbes that are metabolically active but not replicating, and we therefore applied BONCAT (Bioorthogonal Non Canonical Amino Acid Tagging) i.e. a proxy for activity that does not rely on cell division, to measure translationally active cells in soils. We compared the active population of two soil depths from Oak Ridge (TN) incubated under the same conditions for up to seven days. Depending on the soil, a maximum of 25 – 70% of the cells were active, accounting for 3-4 million cells per gram of soil type, which is an order of magnitude higher than previous estimates. The BONCAT positive cell fraction was recovered by fluorescence activated cell sorting (FACS) and identified by 16S rDNA amplicon sequencing. The diversity of the active fraction was a selected subset of the bulk soil community. Excitingly, some of the same members of the community were recruited at both depths independently from their abundance rank. On average, 86% of sequence reads recovered from the active community shared >97% sequence similarity with cultured isolates from the field site. Our observations are in line with a recent report that, of the few taxa that are both abundant and ubiquitous in soil, 45% are also cultured – and indeed some of these ubiquitous microorganisms were found to be translationally active. The use of BONCAT on soil microbiomes provides evidence that a large portion of the soil microbes can be active simultaneously. We conclude that BONCAT coupled to FACS and sequencing is effective for interrogating the active fraction of soil microbiomes *in situ* and provides new perspectives to link metabolic capacity to overall soil ecological traits and processes.

## Introduction

Soil communities are composed of thousands of species that assemble populations of millions to billions of cells within each gram of material ^1,2^. Together, they perform key nutrient cycling functions that as a collective are dominant contributors to Earth’s biogeochemical cycles^3^. Next generation sequencing techniques have enabled the detailed description of the microbial taxa inhabiting soils ^4^, and their comparison across large sets of samples with the aim of pinpointing the drivers of the microbial diversity ^3,5,6^. This enables measurement of the patterns of diversity that emerge in soils, especially in terms of correlation with edaphic factors, such as pH ^7^, soil texture ^8^ or moisture content ^9^, or biological factors, such as species-species interactions, life strategy ^10^ or rank abundance ^6^. Yet, it is challenging to extrapolate from microbial abundance to function, and it is impossible to scale-up these observations to the macroscale^11^. For instance, it is still challenging to predict quantitatively the impact that temperature change will have on decomposition and remobilization of soil organic matter^12^. In order to better couple soil function to microbial activity, one would have to take into account that (i) a large fraction (∼40%) of the microbial diversity retrieved in soils by molecular methods might come from physiologically compromised cells, free DNA ^13^ or dormant cells ^14^ and (ii) microbial activity in the soil will be constrained by geochemical heterogeneities that vary at the microscale^15^. The development of complementary technologies that probe active cells *in situ* will therefore provide an avenue to better link microbial diversity to soil microbial activities ^16^.

Probing active microorganisms *in situ* has been traditionally achieved by using stable isotope probing (SIP) or bromodeoxyuridine (BrdU) labeling. SIP encompasses a series of methods that involve the incorporation of heavy isotopes into newly synthetized DNA^17^ and its separation on a density gradient 18. SIP using labeled ^13^C compounds has shed light onto how the soil microbiome metabolizes certain molecules of interest such as cellulose ^19^ and was also used to track newly formed cells by labeling them with H_2_^18^O ^20^. More recently, BrdU DNA immunocpaturing was implemented in soils, BrdU is a thymidine analog that becomes incorporated into DNA in cells undergoing replication, enabling DNA immunocapturing using BrdU antibodies ^21,22,23^. These methods coupled to high throughput sequencing have enabled the examination of active microbes; however they only capture cells that have undergone division while many cells in soils may be metabolically active yet not replicating. Importantly these methods quantify the amount of labelled DNA rather than the number of active cells in the soil, which can lead to an underestimation of the active fraction given that a large fraction of soil DNA might be relic DNA from physiologically impaired cells^13^.

Other techniques, mostly based on the so called “live/dead” fluorescent DNA stain, attempted to quantify the fraction of active cells in soil through direct counts and found an average of 1.9% cells to be active^24^. In a situation where cells in soil are ∼10 microns apart from each other on average ^25^, and only 1.9% of them are active, it is difficult to imagine how these cells would control carbon dynamics ^26^, and establish complex metabolic networks capable of coordinated responses to changes of conditions^10,27,28^. We hypothesized that at least at some point in time, more cells should be active in order to maintain microbial diversity^29^ and to explain the dynamic of taxa abundance and respiration fluctuation measured in soil^28^. We further expected that some cells may be metabolically active but slowly replicating and that these could potentially be detected through labeling biomolecules that have a faster turnover than the genomic DNA, such as proteins^16^.

Recently, Bioorthogonal Non Canonical Amino Acid Tagging (BONCAT) was reported as an approach to characterizing the active fraction of marine sediment communities^30,31^. This approach uses a relatively fast procedure and small amounts of material, attributes that make it an appealing experimental procedure. This technique consists of incubating the sample with homopropargylglycine (HPG), a water soluble analog of methionine, containing an alkyne group, which is incorporated into newly synthesized proteins^32,33^. Fluorescent dyes are then conjugated to HPG-containing proteins using an azide-alkyne “click chemistry” reaction^32^. As a consequence, cells that were translationally active during the incubation are fluorescently labeled and can be specifically recovered using fluorescence activated cell sorting (FACS) ^34^. BONCAT labels newly synthesized proteins and therefore does not rely on cell division and DNA synthesis to occur, facilitating short term incubations (minutes to hours) and interrogation of slow dividing cells. The sequencing of BONCAT-labeled, FACS-recovered cells could thus be a convenient method to provide a snapshot of the active portion of a soil microbiome.

Here, to our knowledge, we report the first use of BONCAT probing of active members of the soil microbiome, as well as the integration of BONCAT with FACS cell sorting and sequencing of the active cells. For these studies, we incubated undisturbed soils from the Oak Ridge Field Research Site (ORFRS) site with HPG, and sorted labeled cells using FACS. The composition of the active community was determined through 16S ribosomal gene amplicon sequencing. The results were compared with the composition of the total soil microbiome as well as to the ∼700 isolates collected from the same field site. These analyses reveal that a large fraction of the microbiome was translationally active—as high as 25– 70% of cells, which is an order of magnitude higher than previous estimates. Further, a large fraction of the active cells whose taxonomy was resolved were close phylotypes of existing isolates and of major soil taxa identified in a recent global soil survey^35^.

## Results and Discussion

### HPG is actively incorporated by cells *in situ*

We evaluated the utility of BONCAT for identifying translationally active cells from soil systems consisting of a highly heterogeneous matrix, and for the first time. We coupled BONCAT with FACS to detect and recover individual active cells, as opposed to microbial aggregates consisting of hundreds of cells^34^, in order to only sequence the active members of the community. Soil samples were collected at the Oak Ridge Field Research Site (ORFRS) in Oak Ridge, TN, USA and were horizontally cored at 30 cm and 76 cm below surface for the analyses of two distinct communities (Figure 1A). Both depths had low organic carbon (∼0.15%), and nitrogen (<0.05 %) content, and no detectable amount of phosphorus (Figure S1). The 30 cm soil had more quartz and less mica than the 76 cm sample that was composed of more clay. None of these samples had detectable amount of methionine based on LC-MS and therefore it is unlikely that there was significant competition for incorporation of the 50 µM non-canonical methionine (HPG) (Figure S2).

**Figure 1:**
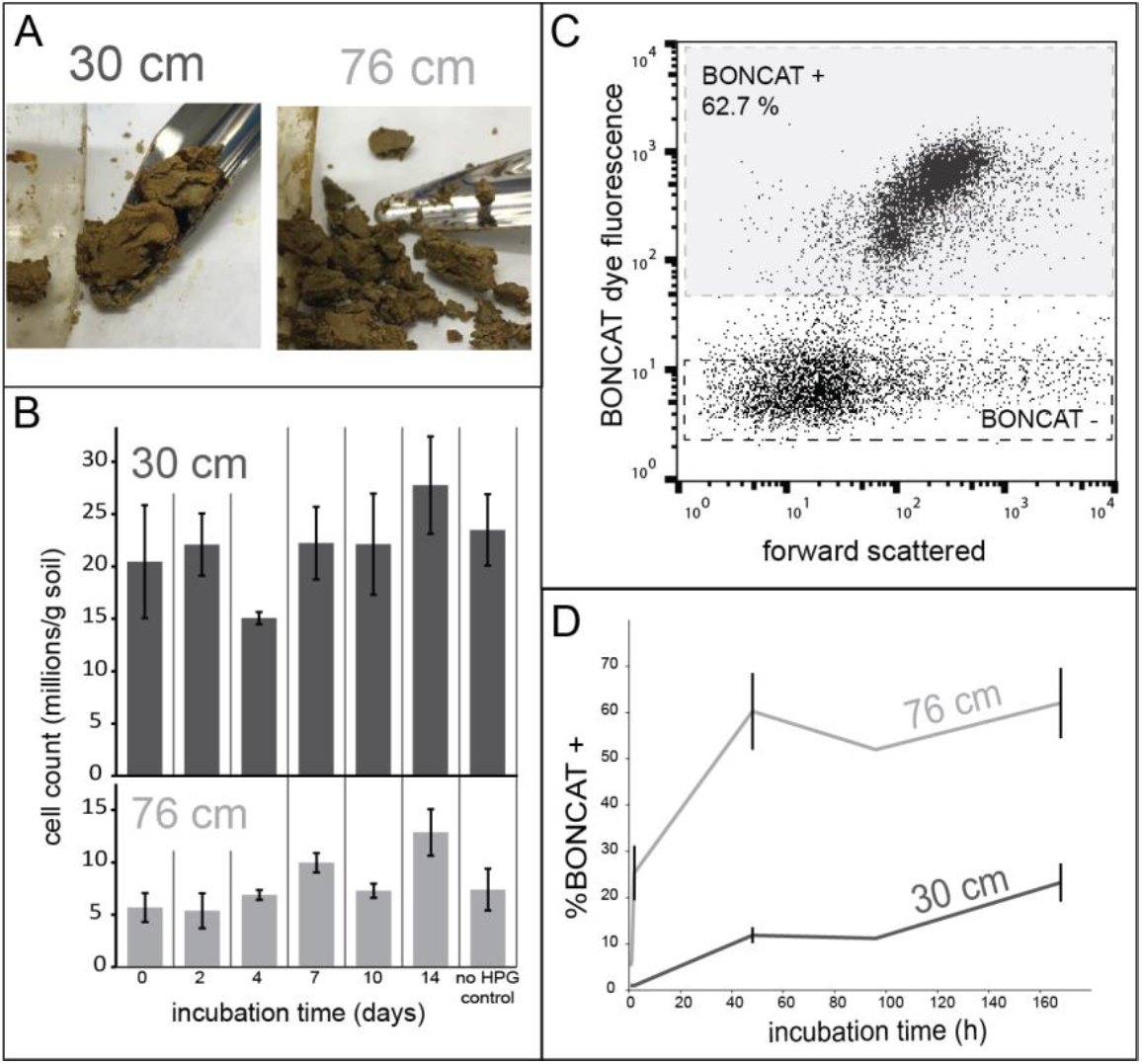
Performing BONCAT-FACS-Seq on soil samples. **(A)** Pictures of the 30 cm and 76 cm soils sampled from the horizontal cores. **(B)** Cell counts over time showing ∼20 million cells per gram at 30 cm and ∼5 million cells per gram at 76 cm. **(C)** Example of BONCAT+ cells as assessed by by flow cytometry. Each dot represents a SYTO59 (DNA dye) positive signal, but only the top population (BONCAT +, grey box) incorporated HPG during incubations. **(D)** Temporal dynamics of BONCAT+ labeling for the 30 cm and the 76 cm sample. Error bars represent standard deviation (n=3).

We first confirmed that HPG was actively incorporated by the cells *in situ*. To do so, we performed a killed control experiment on the 76 cm soil with duplicate samples for each treatment condition. Cells were either fixed before or after incubation with HPG. Cells fixed prior to HPG incubation were not labeled by the BONCAT azide dye, nor were the unfixed cells incubated without HPG. In contrast, both unfixed cells and cells fixed after HPG incubation acquired a distinct green fluorescence signal corresponding to the BONCAT dye. The fraction of fluorescent cells and the per-cell fluorescence intensities were similar between unfixed and post-incubation fixed cells (Figure S3). This confirmed that HPG was only incorporated by active cells, and that fixation was not required for the cycloaddition of the BONCAT azide fluorescent dye. Therefore, all additional experiments were performed with unfixed cells.

### Bias introduced by capturing the cells on a filter as part of the BONCAT procedure

Although the HPG incubation was performed on a minimally disturbed soil sample (directly transferred from the soil core), the cells needed to be detached from the soil matrix and captured on a 0.2 µm filter (see methods) for the click reaction, and subsequently detached again for FACS analysis. We evaluated the bias introduced by these two steps on the microbial community structure, as some cells might detach preferentially from the soil as compared to others being more firmly attached, and the fraction of cells smaller than 0.2 µm^36^ might be lost in the filtration step. For this, we compared the microbial communities retrieved from total soil to the communities captured on the filter.

The microbial community structure retrieved from the total DNA of the 30 cm and 76 cm soil differed at the phylum level (Figure 2A). For example, the 30 cm soil was dominated by Acidobacteria as well as candidate phlya AD3 and GAL15. The 76 cm soil was largely dominated by Proteobacteria with a higher fraction of Bacteroidetes than found in 30 cm soil. At the exact variant sequence (ESV) level, the most abundant ESV at both depths was an Alphaproteobacterium genus *Aquamicrobium* that accounted for 8.77% and 72.9% of the analyzed sequences from 30 cm full soil and 76 cm full soil, respectively. This OTU was only partially captured on the 0.2 µm filters (it represented 2.4% and 3.1% respectively for the 76 cm and the 30 cm filters). The number of OTUs (ESVs clustered at 97% similarity) captured on the filters for the 30 cm sample was half the number retrieved from the total soil, whereas the number captured at 76 cm filters captured greater than or equal to the full soil sample in average (Table S1). The OTUs that were present on the 76 cm filter sample and not retrieved in the soil were found in low abundance (< 0.1 %) and might have come from the rare members of the soil microbiome. Since we observed that the filtering (which was used for the BONCAT cells) introduced a bias, we used the total cells captured on a filter for further comparison with the BONCAT sorted fractions.

**Figure 2:**
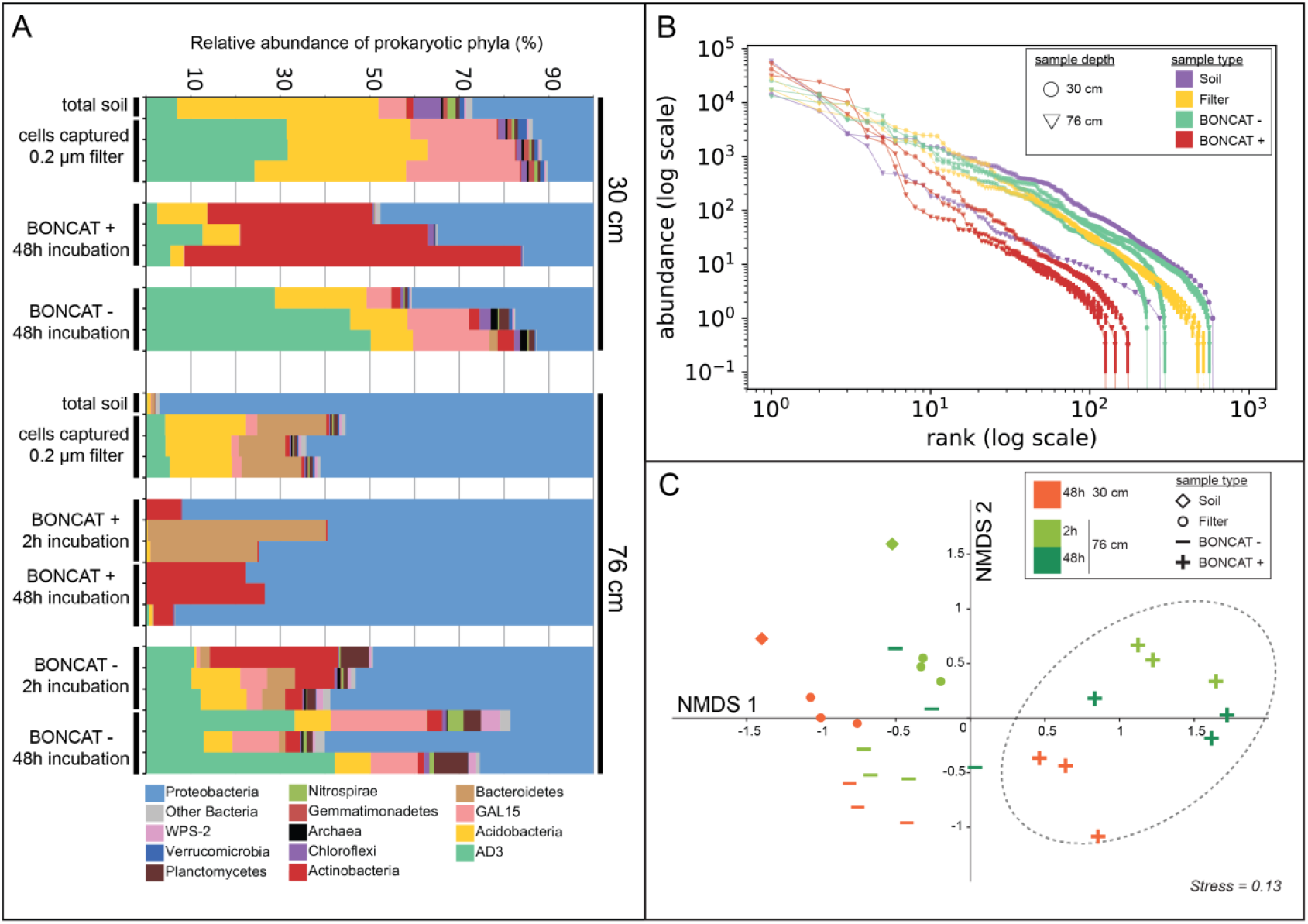
Composition of total community and BONCAT+ cells determined by 16S sequencing. **(A)** Microbial diversity displayed at the phylum level for all samples analyzed. **(B)** Rank *vs.* abundance plot in log-log scale of the libraries of averaged biological replicates, standard deviations are displayed as error bars (n=3). **(C)** NMDS ordination of the Bray Curtis pairwise distance of all libraries. 95% confidence ellipse is displayed on the BONCAT+ group of samples. “Soil” samples are libraries constructed from total DNA extracted from soil, “Filter” samples are DNA extracted from all cells detached from soil and captured on a 0.2 µm filter, BONCAT+ and BONCAT-libraries were constructed from corresponding cell sorted samples.

### BONCAT labeling challenges our view of the active fraction of soil microbes

To identify individual active cells within soils, samples from 30 cm and 76 cm were incubated with HPG and sampled at various time points up to one week (168 h), followed by fluorescent labeling. The total number of cells per gram of soil was stable throughout the incubation period, with ∼20 million cells g^-1^ soil at 30 cm and ∼5 million cells g^-1^ soil at 76 cm (Figure 1B), indicating there was neither acute toxicity leading to massive cell loss nor a stimulation leading to a population bloom during the incubations. While total cell numbers held steady, the fraction of BONCAT+ (Figure 1C) cells increased over time in both soil samples (Figure S4), with a distinct rate of labeling and fraction of labeled cells detected for both soil samples. For example, cells from the 76 cm soil were labeled quickly (clear BONCAT+ population were visible as early as 30 min after incubation) and ∼60% of all cells were labeled by 48 h, whereas cells from the 30 cm soil were labeled more slowly (no BONCAT labeling after 1 h) and only ∼20% total cells were labeled after 48 h (Figure 1D). These differences, which were consistent among biological replicates, suggest that the microbial community found at 76 cm was composed primarily of active cells, while the community at 30 cm had a larger fraction of inactive cells.

Soil activity has traditionally been assessed via bulk measurement of microbial processes (such as CO_2_ evolution or enzymatic activities)^37^, however, both the realization that bulk measurements where poor predictors of soil processes^11^ and that the soil organic matter (including the recalcitrant fraction) is composed of small organic molecules of microbial origin^38^ motivate our examination of active soil microbes at the single cell level. Previous attempts have reported that only a small fraction of cells (0.1 – 2%) are active at once^14^, shaping the view that most soils microbes are dormant and constitute a ‘seed bank’ whose members can become active under favorable conditions^29^. These observations along with the realization that the percent of soil surface area covered by microorganisms might be as low as 10^-6^ % ^39^, and that the localization of soil bacteria *in-situ* still eludes us ^40^, contradict the intuition that a large fraction of the cells should be active to account for the observed bulk activities. The soils that we analyzed here showed that 20 % or more cells were active at both depths, and that this value can be reached within 30 min of incubation, as shown for the 76 cm soil. The high number of active cells we found is an order of magnitude higher than previous estimates, and might be partially explained by the fact that it is a fraction of the soil intact cell detached from the soil, and not of the total cells. Nevertheless, our study shows that there are more active cells than previously thought per gram of soil, a fact that supports the sequences-based network approaches to interrogate soil microbiomes. If a large fraction of cells are co-active it is more likely that they will be able to interact metabolically.

### The BONCAT+ fraction forms a selected sub-set of the total community

To determine the identity of active cells, we sequenced the 16S ribosomal marker genes of BONCAT+ cells from 30 cm and 76 cm soil samples. Specifically, triplicate collections of BONCAT+ cells recovered by FACS (2 h incubation of the 76 cm sample and 48 h incubations of the 76 cm and 30 cm samples) were characterized using iTag sequencing (Table S1). Both soils were sequenced for the 48 h time point, as it represents the beginning of the plateau phase of the BONCAT labeling for both cores (Figure 1D). For the 76 cm soil, the 2 h time point was also sequenced to identify early responders. Unlabeled cells (BONCAT-) were also sorted and sequenced from these time points (Figure S1). In order to compare the BONCAT sorted fractions to the total community at a large scale, we plotted the rank vs. abundance of all libraries (Figure 2B). This plot clearly shows that the BONCAT+ populations separate from the rest of the samples, with a steeper slope reflecting a faster drop of diversity at higher ranks. The pattern for BONCAT-samples was similar to the total community captured on a 0.2 µm filter. In order to assess if this difference was from compositional variation, a beta-diversity metric (Bray-Curtis distance) was computed and ordinated with pairwise distances between samples (Figure 2C). The resulting NMDS plot revealed that all the BONCAT+ sorted fractions from both the 30 cm and the 76 cm formed a distinct group from the rest of the samples (Adonis, F = 2.65, *p* value = 0.001). These results indicate that the pools of BONCAT-cells, although of lower diversity compared to the control total soil and filter samples, were a random subset of the total communities’ analyzed, while the BONCAT+ fraction was clearly composed of a distinct and reproducible subset of the community.

Analyzing the phylogeny of the BONCAT+ samples at the phylum level, we found that at 30 cm, the active fraction was dominated by Actinobacteria (Figure 2A), with one *Arthrobacter* OTU encompassing ∼51% of the retrieved sequences on average (“h” Figure 3A), while the 76 cm active population was dominated by Proteobacteria. At the OTU level (ESVs clustered at 97% similarity) (Figure 3), the BONCAT responders OTU h-e-f were highly active at both 30 cm and 76 cm independent of their abundance in the parent population. For instance, OTU h *Arthrobacter* was only recovered at low abundance (rank 214) from the total cells captured on a filter, while it is the most abundant OTU in the BONCAT+ fraction for this sample. This observation suggests that the activity of these very prominent responders was driven more by the incubation condition than by their rank in the parent community. By contrast, the most abundant members of the 76 cm community (OTU a Figure 3) were BONCAT–, indicating that they did not respond to this incubation condition.

**Figure 3:**
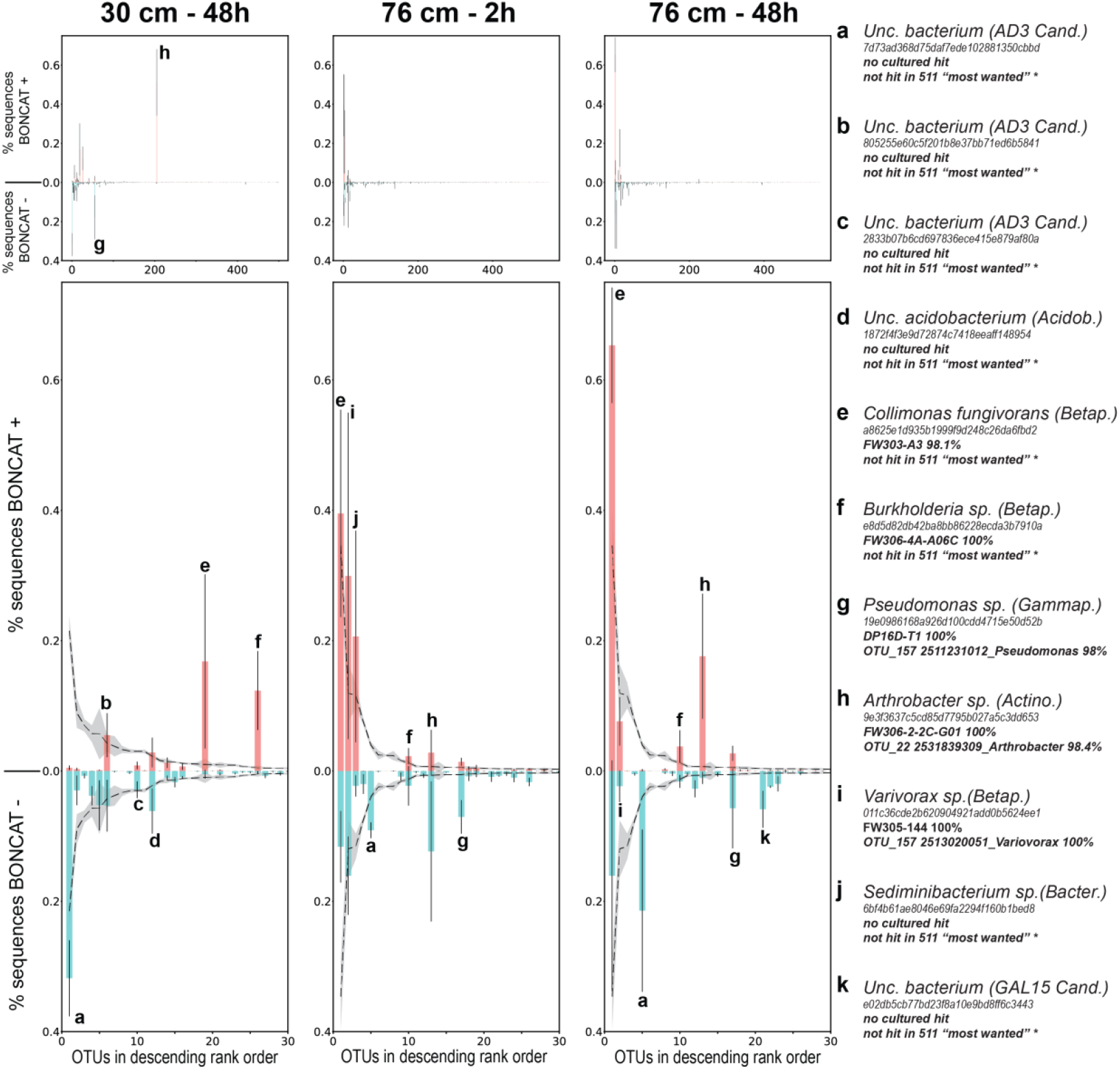
Comparing the composition of the BONCAT+ and BONCAT – populations. **(A)** Relative abundance (in percent, ±SD, n=3) of OTUs present in the BONCAT+ (red) and BONCAT – (blue) for the 30 cm - 48h incubation (left panel), 76 cm - 2 h incubation (middle panel) and 76cm - 48h incubation (right panel). The OTUs have been ranked in descending order from left to right according to their relative abundance on the filter samples (all cells detached and captured on a filter). **(B)** Close-up on the 30 most abundant OTUs overlaid with their abundance on the filter samples (dashed line, ±SD shows as gray shading). The most abundant OTUs are indexed from a to k. Their taxonomy, ID, hit in the ENIGMA culture collection and matches to the 511 most abundant soil microbiome ^35^ is provided on the right legend panel.

Although we are confident that the BONCAT + fraction is composed of translationally active cells, the relative proportion of the OTUs within each library is to be interpreted with some caution, because of factors including potential biases from PCR when producing the iTags libraries^41,42^, and sorting (in the detachment from the filter and DNA staining steps) that impacted estimates of relative abundance. More precisely, a few OTUs account for the majority of the sequences retrieved in the BONCAT fraction, while their abundance were lower in the total community. Given that the size of the total population did not vary during the incubation, this can be explained by two non-exclusive hypotheses: (i) some technical bias in determining relative abundance or (ii) real growth of certain OTUs perfectly balanced by loss of other members. While our experimental design does not allow us to distinguish between microbes that were already active and the ones that were activated by the incubation, it seems reasonable to assume that the signal we measured is a mix of both types.

Another interesting finding from this study is that the BONCAT+ signal plateaued at around ∼4 million active cells per gram of soil, independent of the size of the total population. This raises an interesting hypothesis of a resource limit within these samples that controlled the total number of active cells in each soil sample. To explain the persistence of the inactive members of the community, especially the abundant ones (such as OTU a which is the most abundant OTU at 30 cm), we suggest that they must thrive under other sets of parameters that they encounter in the field but not in our experimental setup. In addition, it is possible that some of the BONCAT- are false negatives due to their inability to incorporate HPG. At this point, it is difficult to estimate the bias introduced by the preferential incorporation of HPG to certain species compared to others, but the fact that BONCAT+ cells belonged to 251 different OTUs spanning 17 bacterial phyla and accounted for up to ∼70% of the detachable cells, suggests, as previously noted ^34^, that HPG is in fact incorporated by a large set of bacterial species.

We see that BONCAT labeling probes a variety of organisms, many of which are not cultivated. Therefore, BONCAT could also serve the purpose of pinpointing relevant culture conditions for to date uncultivated microbes to be active, and from which they might be isolated more easily than from standard laboratory conditions. For instance, the AD3 candidate division was first proposed in 2003 from a study of sandy soils ^43^ and was repeatedly found in soil since then. However, there is still no cultivated representative of this phylum reported. We found activity for some AD3 members under our selected incubation conditions, which thus provide a new starting point for cultivation efforts for this phylum.

### BONCAT responders closely related to cultured and generally abundant soil organisms

We further asked how the culture collection available for this experimental site captured the diversity of the active, and presumably ecologically relevant, fraction of the community. For this, we compared 16S sequences of BONCAT+ cells and total cells libraries (both total soil and cells captured on a filter) to 16S sequences from 687 isolates collected from this same location. Surprisingly, 77% to 98% of total sequences from BONCAT+ cells shared >97% sequence similarity with an isolate collected from the same location—consistent with the view that the active microbes lend themselves to isolation. This is despite the fact that when looking at the total community of cells (filter sample), only 7% and 2% of the total sequence reads from the 76 cm and the 30 cm respectively shared >97% sequence similarity with the isolates, suggesting the isolated OTUs were not part of the dominant members of the community (Figure 4A).

**Figure 4:**
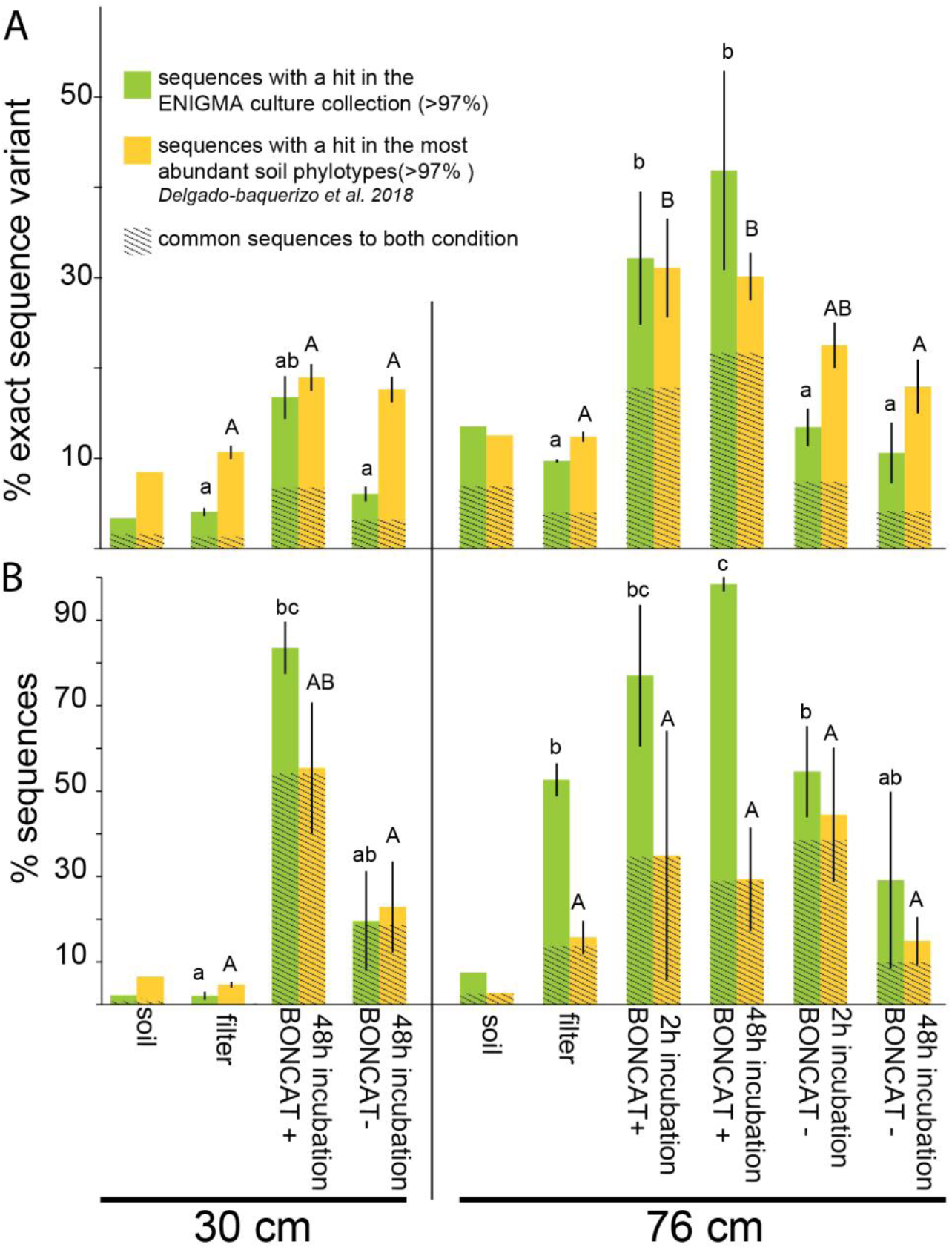
Prevalence of soil isolates and ubiquitous soil OTUs among BONCAT+ cells. **(A)** Percent ESVs and **(B)** percent sequences from the current libraries with a hit (>97% sequence similarity) in the ENIGMA culture collection (this collection contains 697 full-length 16S rDNA from strains that were isolated from the same field site as the samples considered in this study) (green), in the set of 511 phylotypes identified as the most abundant by ^35^ (yellow) or both (dashed area). Data are average (n=3) ±SD, letters indicate ANOVA post-hoc significant differences. “soil” samples are libraries constructed from total DNA extracted from soil, “filter” samples are DNA extracted from all cells detached from soil and captured on a 0.2 µm filter, BONCAT+ and BONCAT-libraries were constructed from corresponding cell sorted samples.

It is interesting that in our incubation condition, abundant OTUs (e, f, h and i, Figure 3) with close cultured representatives were translationally active in an oligotrophic soil (∼0.15% TOC) without any addition of nutrients other than HPG. Importantly, the cell counts remained stable throughout the incubation (Figure 1B), suggesting that the predominance of these OTUs in the BONCAT+ fraction did not result from them overgrowing the community during the incubation. Soil microbiomes are typically highly diverse and composed of largely uncultivated lineages^3^. Our data suggests that at least in this case, cultivated microbes comprise a substantial portion of the active cells. Although unexpected, this observation aligns well with the recently published contribution from Delgado-Baquerizo *et al.* 2018 ^35^ that identified a list of 511 phylotypes (OTUs with 97% cutoff) encompassing 44% of the microbial diversity of soils worldwide. Among these phylotypes, 45% had a cultured representative, suggesting that only a small number of microbial phylotypes might be globally relevant for soil microbiomes, and that cultivation efforts have already yielded to isolate representatives of a substantial amount. In order to further compare our dataset with these 511 ubiquitous soil phylotypes, we ran BLAST comparisons on a set of representative sequences of our libraries OTUs and recovered the >97% hits (Figure 4A and 4B). We found that there was an overlap between the sequences found in the ENIGMA culture collection and from the 511 reference phylotypes. Three of the most abundant BONCAT+ OTUs retrieved belonged to the 511 prominent members of the global atlas for soil microbiome^35^ (e.g. OTU g, h, i Figure 3).

### Exploring the link between abundance and activity in soils

Most if not all structures of microbial communities follow a power low rank-abundance trend with a few highly abundant members, and a large number of rare members, which we also see for our data (Fig 2B). In order to persist in a community, all members should have a positive growth over death population ratio. However, in order to become abundant, a particular member must have higher growth rates and/or lower mortality rates than other members. Therefore, the makeup of the total community is an integration over time of microbial populations’ turnover. It was proposed that dormant microbes play an important role in maintaining microbial diversity, acting as a “seed bank” where different OTUs become active under favorable conditions^29^. As a corollary it is thought that most activity is concentrated in hotspots with high organic input such as the rhizosphere^44^. Our data suggest that under a given set of conditions, only a small subset of the soil microbiome OTUs are active (88 OTUs on average in this study, Table S1), however these account for a large part of the cell population (20 % - 60 %) even in areas of soils that likely are not hotspots.

Despite dramatic differences in their initial abundance, a subset of consistently active cells is present across samples (e.g. OTUs e, f, h in Figure 3). This last observation is reminiscent of the “scout” model ^45^: In this model, most cells are not actively growing, and ‘scout’ cells randomly “awaken” and start to divide if the resource conditions are favorable to do so. If the “scout” model holds true, one would indeed expect that under a given set of incubation conditions (as provided here), only a small fraction of the microbial diversity would wake up and an even smaller one would be able to thrive. We see examples of OTUs that appear to follow this ‘awakening’ behavior: for example, OTU h (Figure 3) was present at a very low abundance at 30 cm, and became the most abundant OTU in the BONCAT+ fraction after 48 h. As a corollary of the ‘scout’ model, because of spontaneous awakening, it is expected that active and inactive cells coexist for a given species, a prediction that is also verified by our data (e.g. OTU b and d Figure 3). These observations are also consistent with previous studies that have identified that even non-sporulating bacteria exhibit periods of transient dormancy^46,47^ and that bacterial signaling molecules exist that promote cell resuscitation^48,49^ whose could explain why only certain populations become active.

### Conclusions

We find BONCAT to be a useful tool for analysis of the active fraction of soil microbiomes when coupled to fluorescence activated single cell sorting (FACS) and sequencing. BONCAT enables separating active cells from free DNA, dormant microbes and physiologically impaired cells. It can be viewed as a filter that focuses environmental DNA analyses on the active and likely ecologically relevant cell fraction in a given environmental condition. As all filters, the BONCAT procedure also introduces biases and will need to be benchmarked against other activity probing strategies. Nonetheless, we showed that BONCAT-FACS-Seq can be used to track the active cell population dynamics and dissect the behavior of active members at the phylum or OTU level. Our experiments resulted in consistent enrichment of a specific set of organisms in the BONCAT+ fraction that differs from the total community. Surprisingly, we found that a large fraction of the cells were active under our incubation conditions (25 % - 70 %), which contradicts the common view that most soil organisms are inactive (Figure 5). Overall, our data shows that the application of BONCAT-FACS-Seq is a powerful approach that can provide important new insights into soil microbiomes with the potential to help reconcile functional measurement to microbial diversity. Given these encouraging results and the relative simplicity of the approach we predict that it is going to be widely used in future applied and fundamental soil microbiome research.

**Figure 5:**
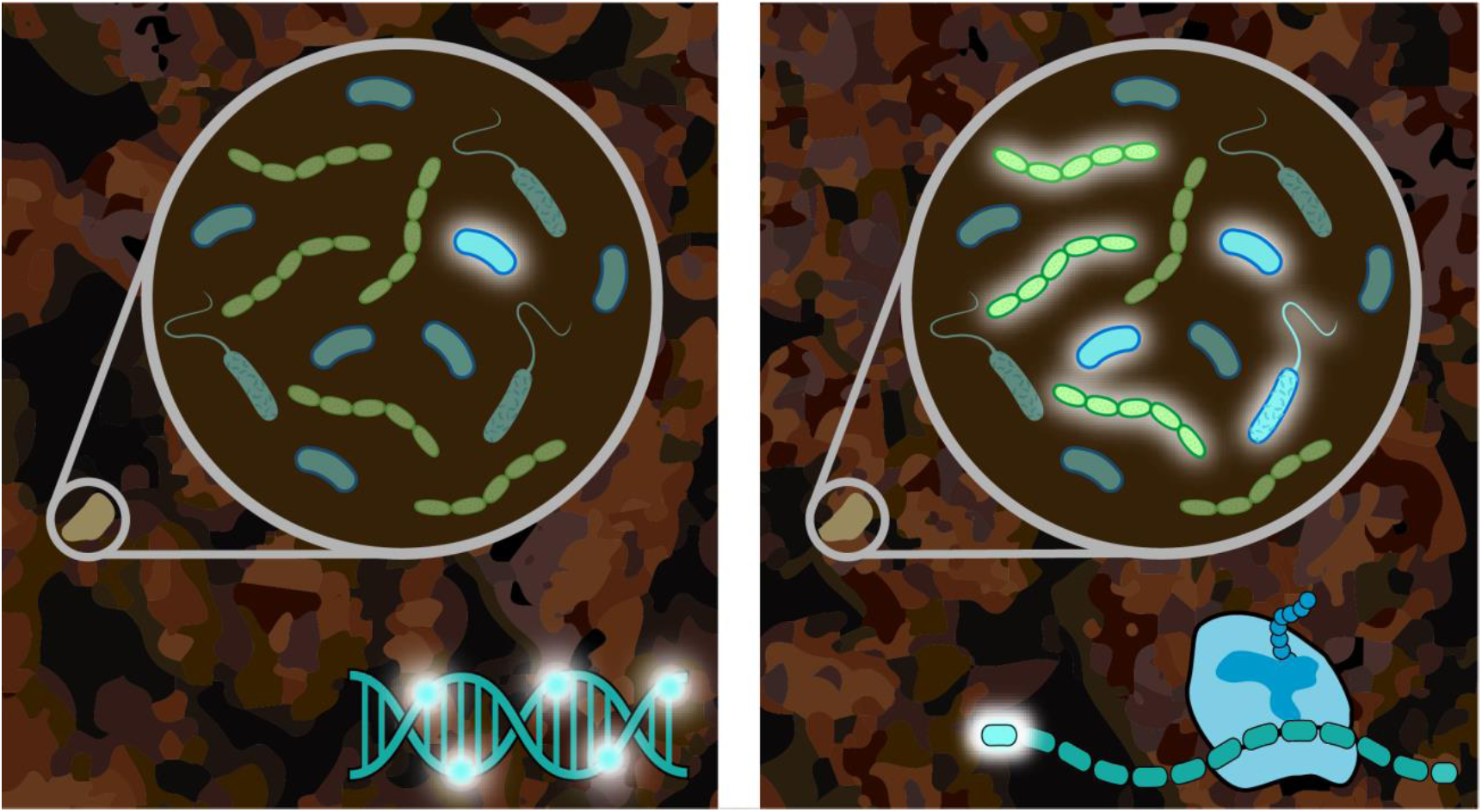
The use of BONCAT is adding a large fraction of active microbe on the soil microbiome picture (Left panel) Traditional view, based on DNA labelling (right bottom corner) showing that 1.9 % of cells on average are active in soils^24^, **(Right panel)** By labeling proteins (right bottom corner) we find a large fraction (up to 70%) of the cells can be active at once *in situ* in our soil samples.

## Material and methods

### Samples collection and incubation condition

Two 4 cm diameter sample soil cores were collected horizontally from Oak Ridge, TN (GPS 35.941133, −84.336504) on January 24th 2017 from a silt loam area. A vertical trench was made and a first core was taken at 30 cm depth while the second one was collected at 76 cm depth. Both cores were shipped cooled and where stored in the dark at 4°C until processing. At the time of the experiment (within 1-3 months after collection) a piece of ∼1 g of soil was sampled from the distal part of the core under sterile conditions for each replicate and placed into a 10 ml culture tube, the full design can be found in Figure S1. Each replicate was incubated with 2 ml of 50 µM L-Homopropargylglycine (HPG, Click Chemistry Tools, Scottsdale, AZ, USA) in sterile water at 15°C in the dark. This temperature was chosen because it is the average surface temperature at the field site. At the end of the incubation period (spanning 0.5 h to 168 h, see Fig. Figure S1) 5 ml of 0.02% Tween^®^ 20 (Sigma-Aldrich, ST Louis, MO, USA) in phosphate saline buffer (1X PBS) was added to each tube and further vortexed at maximum speed for 5 min (Vortex-Genie 2, Scientific Industries, Inc., Bohemia, NY, USA) in order to detach cells from the soil particles. Culture tubes were then centrifuged at 500 g for 55 min (centrifuge 5810R, Eppendorf, Hamburg, Germany) and the supernatant was frozen at −20°C in 10% glycerol (Sigma-Aldrich, ST Louis, MO, USA) until further processing.

### Killed control experiment

We performed a killed control experiment to validate the active incorporation of HPG by the cells; every condition was tested for a biological duplicate. Samples from the 76 cm were either fixed with 3 % paraformaldehyde (PFA, Sigma-Aldrich, ST Louis MO, USA) incubated for 1 h at RT prior or after incubation with HPG. These samples were compared to no HPG control (with and without PFA) and non-fixed samples. Incubation times were 2 h and 48 h. This set of sample was handled as previously described, cells were detached from the soil and frozen stock in 10 % glycerol were kept at −20 °C until further evaluation of HPG incorporation, see below.

### Soil properties, mineral and organic composition of the soils

Bulk X-ray powder diffraction was used to analyze the mineralogical composition of the soils cores. Powdered samples were loaded on an autosampler in a Rigaku SmartLab X-ray diffractometer (Rigaku, The Woodlands, TX, USA), using a Bragg-Brentano geometry in a theta-theta configuration. Data were collected from 4° to 70° of 2θ, using Cu Kα radiation. After manual identification of the phases present, a Rietveld refinement was performed to obtain their weight fractions, using the software MAUD ^50^.

### Click reaction - BONCAT stain

A volume of 700 µl of frozen cells of each sample were allowed to thaw at 4°C for ∼1 h h. In the meantime, the click-reaction mixture was prepared by mixing the dye premix with the reaction buffer. This premix consisted of 5 µl copper sulfate (CuSO_4_ 100 µM final concentration), of 10 µl tris-hydroxypropyltriazolylmethylamine (THPTA, 500 µM final concentration), and of 3.3 µl (FAM picolyl azide dye,5 µM final concentration). The mix was incubated for 3 min in the dark before being mixed with the reaction buffer, which was made of 50 µl sodium ascorbate freshly prepared in 1X PBS at 5 mM final concentration and 50 µl of aminoguanidine HCl freshly prepared in 1X PBS at 5 mM final concentration and 880 µl of 1X PBS. All reagents were purchased from Click Chemistry Tools (Click Chemistry Tools, Scottsdale, AZ, USA). Once thawed, the cells were captured on a 0.2 µm GTTP isopore™ 25 mm diameter filter (MilliporeSigma, Burlington, MA, USA) and rinsed with 7 ml 1X PBS. The filter was then placed on a glass slide and 80 µl of the click reaction mixture was quickly added before covering the filter with a coverslip to avoid excess oxygen during the click reaction. The slides were incubated in the dark for 30 min and each filter was then thoroughly washed three times in a succession of three baths of 20 ml 1X PBS for 5 min each. The filters were finally transferred to 5 ml tubes (BD-Falcon 5 ml round bottom tube with snap cap, Corning^™^, Corning, NY, USA) with 2 ml of 0.02% Tween^®^ 20in PBS, with the cells facing inwards and vortexed at maximum speed for 5 min to detach the cells. The tubes were incubated for 20 min at 25°C, and subsequently stored at 4°C. Before being loaded onto the cell sorter (BD-Influx^™^, BD Biosciences, San Jose, CA, USA), the samples were filtered through a 35 µm filter (BD-falcon 5ml tube with cell strainer cap, Corning^™^, Corning, NY, USA). A water incubated sample was clicked along with each set of samples to define the BONCAT staining background of each single click reaction.

### Flow cytometer, cell count and cell sorting

For the cell counts, the cells were prepared the exact same way as described above, but the click reaction was omitted and the cells detached from the filter were stained 1X SYBR^™^ (ThermoFisher Scientific, Invitrogen, Eugene OR, USA). For the evaluation of the BONCAT stained samples, cells were counterstained with the SYTO^™^ 59 (ThermoFisher Scientific, Invitrogen, Eugene OR, USA) DNA dye for 5 min at RT at 0.5 µM. The cell sorter (BD-Influx^™^, BD Biosciences, San Jose, CA, USA) was setup to capture the FAM picolyl azide dye (excitation = 490 nm/ emission = 510 nm) in the green channel off a 488nm blue laser and the counter DNA stain (excitation = 622 nm, emission = 645 nm) in the red channel off of a 630 nm red laser. A first gate was drawn on the SYTO positive (SYTO+) particles, under the assumption that this would capture the cells. SYTO + events accounted for 0.1 - 5 % of the events depending on the samples, most of the events being abiotic, most probably clays or other minerals. The BONCAT positive (BONCAT +) and BONCAT negative (BONCAT -) where further gated as a sub-fraction of the SYTO+ cells based on the BONCAT dye fluorescence. The water incubated sample was used as a negative control to define the level of nonspecific BONCAT stain fluorescence, the BONCAT - gate was drawn under that line and BONCAT + gate was such that <0.5% of negative control cells were in it. The percent of BONCAT + determined for a timecourse for both the 30 cm and the 76 cm sample and guided the sorting decisions. We decided to sort three biological replicates at two incubation time points for the 76 cm sample (2h and 48h) and three biological replicates at one time point for the 30 cm sample (48h). A total of 35k-75k cells (see table Figure 1B for detailed counts) were sorted in parallel for the BONCAT + and BONCAT - gates into a 96 well plate. Plates were frozen at −80°C until processing.

### Total DNA extraction from soil and filters

In order to compare sorted cells to the soil microbiome, total purified DNA was prepared from the soil cores and the cells captured on a 0.2 GTTP isopore™ 25 mm filter (MilliporeSigma, Burlington, MA, USA), respectively to account for the bias of the first step of the BONCAT process. We used the Qiagen-MoBio Power soil DNA kit (Qiagen, Hilden, Germany) following the manufacturer instructions, except for the lysis step that was performed by shaking the tubes at 30 Hz for 10 min in a tissue homogenizer (TissueLyser II, Qiagen, Hilden, Germany).

### Libraries preparation and sequencing

In order to pellet the sorted cells, the 96 well plates were centrifuged at 7200 x *g* for 60 min at 10°C. The plates were further centrifuged upside-down for 20 s at 60 x *g* to remove supernatant. The pelleted cells were lysed using PrepGEM (zyGEM, Charlottesville, VA, USA) chemical lysis in 2 µl reactions following manufacturer’s recommendation. 0.2 µl of 10X Green buffer, 0.02 µl of PrepGEM, 0.02 µl of lysozyme and 1.8 µl of water were added to each well. Note that six empty wells were submitted to PrepGEM lysis and library construction to account for potential contaminant. The plates were then placed in a thermocycler for 30 min at 37°C and 30 min at 75°C. The iTag PCR was performed directly on the cell lysate following the JGI standard operating protocol (https://jgi.doe.gov/user-program-info/pmo-overview/protocols-sample-preparation-information/). /). Briefly, the V4 region of the 16S rDNA was amplified using the universal primer set 515F (5’-GTGYCAGCMGCCGCGGTAA-3’), 806R (5’-GGACTACNVGGGTWTCTAAT-3’). The adapter sequences, linkers and barcode were on the reverse primer. The 16S PCR was performed in a final volume of 25 µl (10 µl of the 5 Prime master mix, 0.5 µl of the forward primer (at 10 µM), 1.5 µl of the reverse primer (at 3.3 µM), 0.44 µl of BSA, 10.5 µlµl of water and 2 µl of cell lysate). The PCR condition was as follows: after an initial denaturation step at 94°C for 3 min, 30 PCR cycles occurred consisting of a 45 sec denaturation step at 94°C followed by a 1 min annealing step at 50°C and a 1.5 min elongation step at 72°C. A final elongation step of 10 min at 72°C was further added to finish all incomplete target sequences. The V4 region of the 16S rDNA from the total DNA extracted from the soil and from the cells enriched on filters were also amplified using the same PCR conditions. The PCR products were cleaned using the Agencourt AMpure XP beads solution (Beckman Coulter Life Sciences, Indianapolis, IN, USA) to remove excess primers and primer dimers. PCR products were incubated with 80% (v/v) beads for 5 min at 25 °C before being placed on a magnetic holder (MagWell™ Magnetic Separator 96, EdgeBio, San Jose, CA, USA). The supernatant was removed and the beads were washed with 70% v/v ethanol three times before being resuspended in 11 µl of water. The total DNA extracts were processed in parallel, the only difference being that the iTag PCR was performed in 50 µl final volume and the PCR product was resuspended in 16 µl water after the bead clean-up step. PCR products were run on a High Sensitivity DNA assay Bioanalyzer chip (2100 Bioanalyser, Agilent, Santa Clara, CA, USA) to confirm fragment size and concentration. PCR products were pooled to an equimolar concentration and run on the Illumina MiSeq platform (Illumina, San Diego, CA, USA). Sequences data have been archived under the Bioproject ID PRJNA475109 at the NCBI.

### Sequences processing

The sequences were processed using qiime2 v2017.9^51^. The sequences had been demultiplexed by the JGI sequencing platform, after being deinterleaved, they were imported in qiime2 using the *fastq manifest* format. Sequences were further denoised, the primer trimmed (20 nucleotides from each side) and paired using DADA2^52^, as implemented in the qiime *dada2 denoise-paired* plug-in. This step also included a chimera check using the *consensus* method. The output was a table of 4’063 exact sequence variants (ESVs) of 6’419’059 sequences. 130 ESVs had at least one hit in one of the six no template controls and were not considered for further analysis. The filtered table had 6’110’776 sequences gathered into 3’933 ESVs, and the median value was 205’167 sequences per sample. The ESVs were further clustered into operational taxonomic units (OTU) at a threshold of 97% similarity using the *vsearch cluster-features-de-novo* plug-in. The clustered OTU table had 1’533 OTUs in total. The absolute number of OTUs can vary by up to three orders of magnitude depending on the technique used ^53^, DADA2 is known to return a more conservative number than the previously widely used upfront clustering methods by decreasing the number of false positives^52^. This relatively low number is also consistent with the very low level of organics (carbon and nitrogen) in these soils, which total organic carbon (TOC) are comparable to un-colonized arid lands where microbial diversity is reduced^54^. The taxonomy of the representative sequences was assigned using the *feature-classifier classify-sklearn* plug-in (https://data.qiime2.org/2018.2/common/gg-13-8-99-515-806-nb-classifier.qza). This classifier was trained on the Greengenes database 13_8 99% trimmed to the amplified region (V4 515F/806R). If the classifier could not assign the representative sequences at the phylum level, then they were manually checked on the most up-to-date Silva SINA alignment service (https://www.arb-silva.de/aligner/) and the Silva classification was retained. The OTU table with assigned taxonomy was used to build the bar graph at the phylum level and all downstream analyses. Bray Curtis pairwise distance beta-diversity metric was computed on the OTU table and the obtained triangular distance matrix was ordinated using NMDS.

### Comparison with reference dataset

We compared our iTag data with the 697 full-length 16S rDNA of the ENIGMA Project’s existing culture collection from this field site and with the 511 16S rDNA sequences of the most abundant and widespread soil microbiome members, retrieved from Delgaldo-Baquerizo et al. 2018 ^35^. We performed a nucleotide BLAST of one representative sequence per ESV against the ENIGMA isolate database or the “511 most wanted soil phylotypes”^35^ database using Geneious R9^©^. A cutoff of >97% similarity was used to determine if a sequence from our dataset had a match in the ENIGMA isolate database and/or the “511 most wanted soil phylotypes” database.

### LC-MS Soil metabolomics – targeted analysis for soil methionine that may interfere with BONCAT

Triplicates of 2 g of soils from 30 cm and 70 cm were extracted using 8 ml of LCMS grade water and incubated 1 h on an overhead shaker at 4°C. Aqueous extractable components were collected by removal of insoluble material with centrifugation at 3220 g for 15 min at 4°C, filtration of supernatants through a 0.45 µm PVDF syringe filter (MilliporeSigma, Burlington, MA, USA), followed by lyophilization of filtrates to remove water (Labconco 7670521, Kansas City, MO, USA). Dried samples were then resuspended in 500 µl of LCMS grade methanol, bath sonicated at 25 °C for 15 min, and then clarified by filtration through 0.2 µm PVDF microcentrifugal filtration devices (1000 g, 2 min, 25 °C). Methanol extracts were spiked with an internal standard mix (^13^C,^15^N universally labeled amino acids, 767964, Sigma-Aldrich, USA, which included standard amino acids, including methionine, at a final concentration of 10 µM). Metabolites in extracts were chromatographically separated using hydrophilic liquid interaction chromatography on a SeQuant 5 µm, 150 x 2.1 mm, 200 Å zic-HILIC column (1.50454.0001, Millipore) and detected with a Q Exactive Hybrid Quadrupole-Orbitrap Mass Spectrometer equipped with a HESI-II source probe (ThermoFisher Scientific). Chromatographic separations were done by an Agilent 1290 series HPLC system, used with a column temperature at 40 ^°^C, sample storage was set at 4 ^°^C and injection volume at 6 µl. A gradient of mobile phase A (5 mM ammonium acetate in water) and B (5 mM ammonium acetate, 95% v/v acetonitrile in water) was used for metabolite retention and elution as follows: column equilibration at 0.45 ml.min^-1^ in 100% B for 1.5 min, followed by a linear gradient at 0.45 ml.min^-1^ to 35% A over 13.5 min, a linear gradient to 0.6 ml.min^-1^ and to 100% A over 3 min, a hold at 0.6 ml.min^-1^ and 100% A for 5 min followed by a linear gradient to 0.45 ml.min^-1^ and 100% B over 2 min and re-equilibration for an additional 7 min. Each sample was injected twice: once for analysis in positive ion mode and once for analysis in negative ion mode. The mass spectrometer source was set with a sheath gas flow of 55, aux gas flow of 20 and sweep gas flow of 2 (arbitrary units), spray voltage of |±3| kV, and capillary temperature of 400 ^°^C. Ions were detected by the Q Exactive’s data dependent MS2 Top2 method, with the two highest abundance precursory ions (2.0 m/z isolation window, 17,500 resolution, 1e5 AGC target, 2.0 m/z isolation window, stepped normalized collisions energies of 10, 20 and 30 eV) selected from a full MS pre-scan (70-1050 m/z, 70,000 resolution, 3e6 AGC target, 100 ms maximum ion transmission) with dd settings at 1e3 minimum AGC target, charges excluded above |3| and a 10 s dynamic exclusion window. Internal and external standards were included for quality control purposes, with blank injections between every unique sample. QC mix was injected at the start and end of the injection sequence to ensure the stability of the signal through time and consisted of 30 compounds spanning a large range of *m/z*, RT and detectable in both positive and negative mode. Extracted ion chromatograms for internal standard compounds were evaluated using MZmine version 2.26 ^55^ to ensure consistency between injections. Samples were analyzed using Metabolite Atlas ^55^ (https://github.com/biorack/metatlas). Briefly, a retention time corrected compound library generated by linear regression comparison of QC standards against an in house retention time (RT)-m/z-MSMS library of reference compounds analyzed using the same LCMS methods was used for compound identification in samples where measured RT, m/z and fragmentation spectra were compared with library predicted RT, theoretical m/z, library detected adducts and library MSMS fragmentation spectra. Compounds identification were retained when peak intensity was > 1e4, retention time difference from predicted was < 1 min, m/z was less than 20 ppm from theoretical, expected adduct was detected and at least 1 ion fragment matched the library spectra and were more abundant in at least one sample as compared to the average value + 1 SD of the extraction controls. Only 8 compounds met these criteria; average peak heights from the extracted ion chromatograms are reported in Figure S5. The signal was very low overall owing to the low amount of organics in these soils. We checked for the presence of methionine manually using MZmine version 2.26 ^32^ and confirmed that there were no detectable of methionine in any of the sample analyzed. Metabolomics data has been deposited JGI genome portal #1207416 along with the analysis file #1207417.

## Acknowledgment

The authors would like to acknowledge Dominique Joyner for the samples collection and shipping, Marco Voltolini (LBNL) for his support in the X-Ray diffraction analyses, and the JGI sequencing group for its technical assistance. This work was funded by a discovery proposal awarded to T. Northen as part of the ENIGMA (Ecosystems and Networks Integrated with Genes and Molecular Assemblies http://enigma.lbl.gov) Scientific Focus Area Program at Lawrence Berkeley National Laboratory; the work conducted by the U.S. Department of Energy Joint Genome Institute, a DOE Office of Science User Facility, both supported by the Genomic Sciences Program, Office of Biological and Environmental Research, in the Office of Science of the U.S. Department of Energy under Contract No. DE-AC02-05CH11231.

---

**Figure S1:**
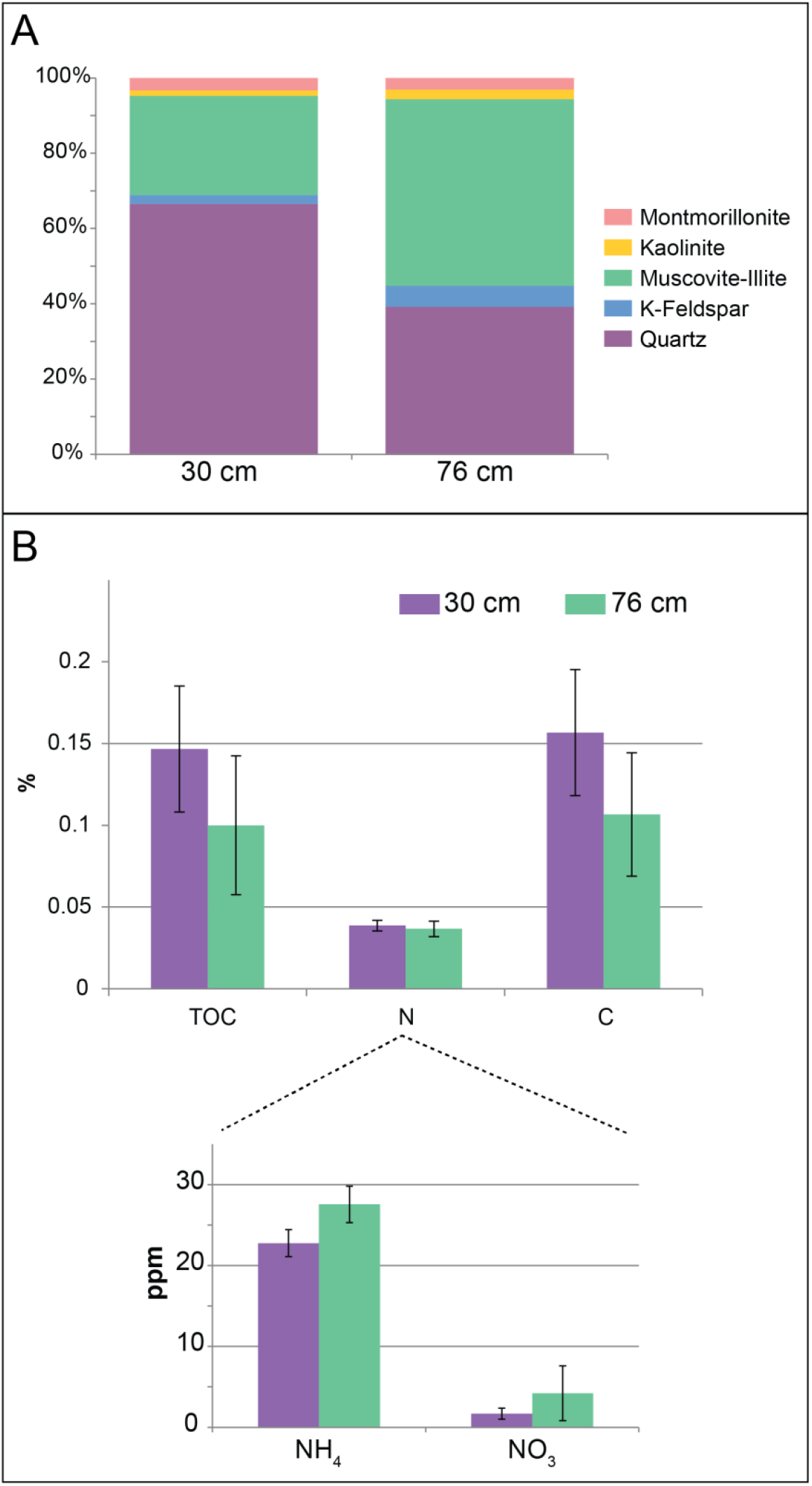
Soils properties. **(A)** Mineral composition **(B)** Total C, TOC, Total N, ammonium and nitrate concentration (n=3). Ppm, parts per million.

**Figure S2:**
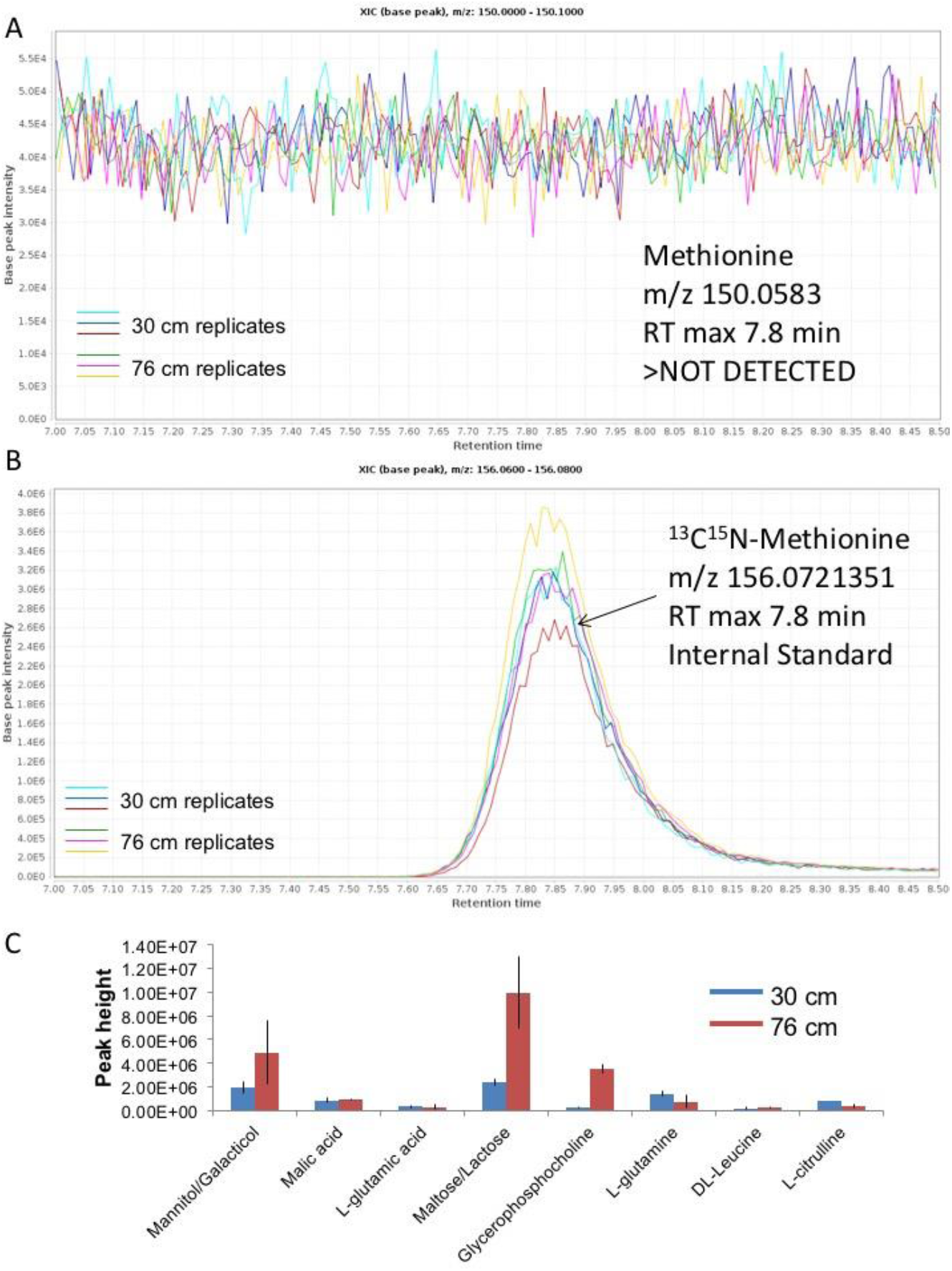
LC-MS analysis of the full soil water extract showing that **(A)** there is no detectable methionine, although **(B)** the spiked labeled heavy methionine was easily detected in all samples. **(C)** Peak height of the 8 compounds that passed our identification criteria (see methods). Data are means ± SD (n=3).

**Figure S3:**
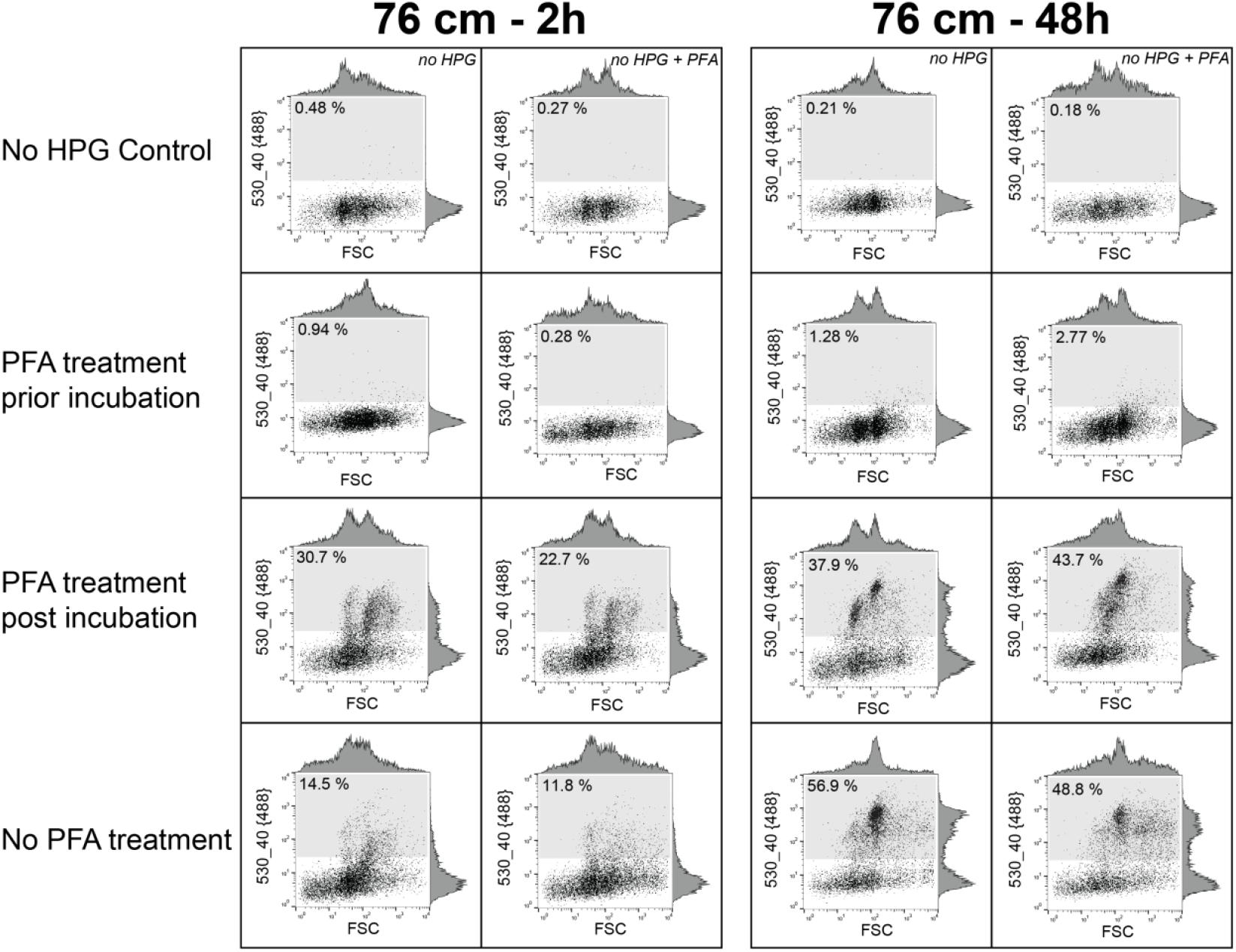
BONCAT labeling of fixed and unfixed cells. Cells stained with SYTO dye plotted according to their forward scatter signal (FSC, x-axis) and BONCAT fluorescence, y-axis) in log-log scale. The distribution of the events along the x and y-axis is shown respectively on the density plot on the top and on the right of each graph. The BONCAT+ cells gate is displayed as a gray box in each plot, and the percent cells in the BONCAT+ gate is indicated in the top left corner of the box. The left two columns are biological replicates from 76 cm soil incubated for 2 h, and the right two for 48 h. Each row of panels corresponds to a different treatment, the first row being control samples without HPG (with or without PFA fixation), the second corresponds to samples that were pre-treated with PFA before incubation, while the third corresponds to samples were cells were fixed with PFA after incubation. The last row corresponds samples that were not fixed, *i.e* the same treatment used for when sorting and sequencing BONCAT+ and BONCAT– cells (Figure S3).

**Figure S4:**
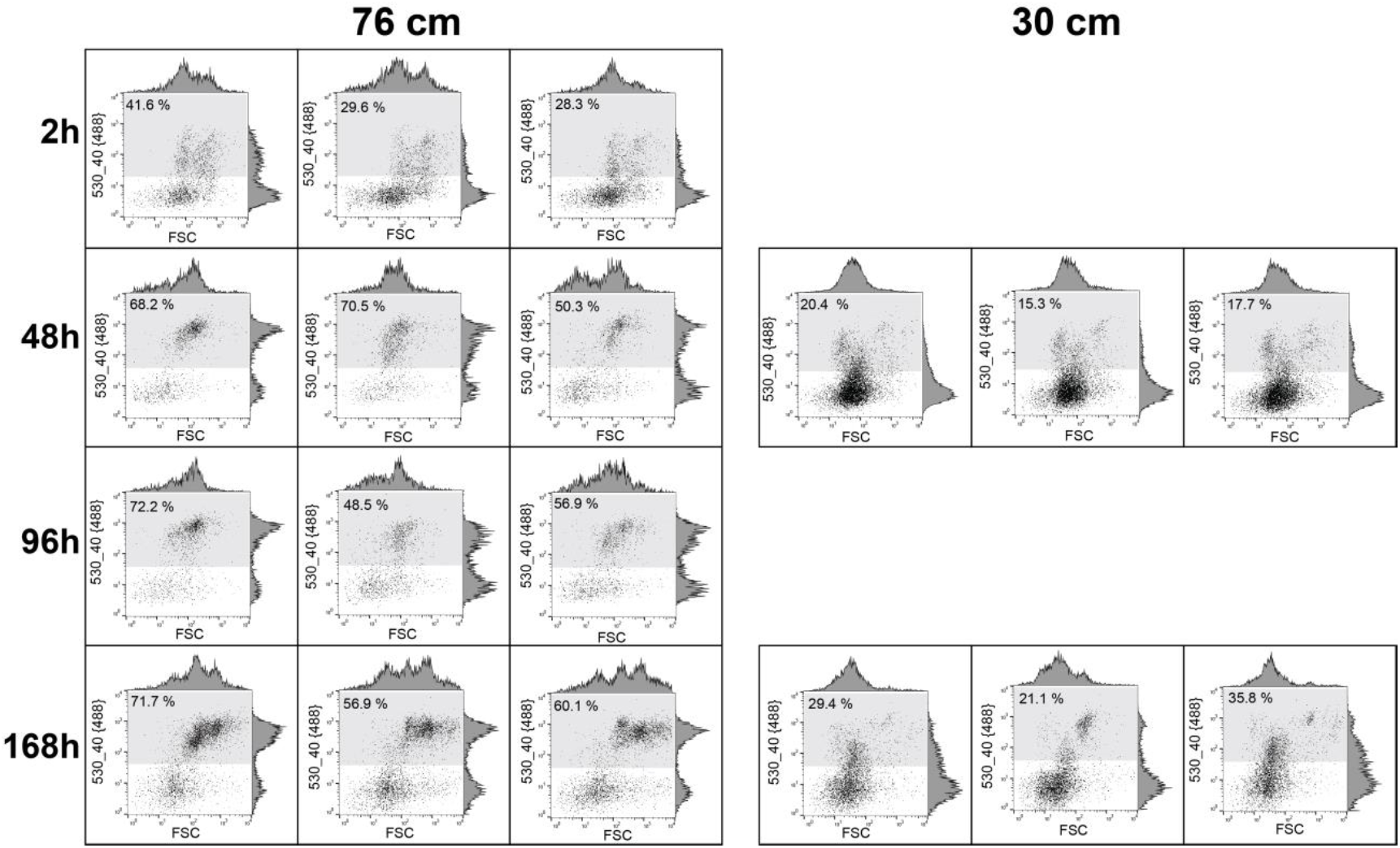
Evaluation of BONCAT+ size fraction. Cells stained with SYTO dye plotted according to their forward scatter signal (FSC, x-axis) and BONCAT fluorescence (Ex: 488nm/Em: 530nm, y-axis) in log-log scale. The distribution of the events along the x and y-axis is shown respectively on the density plot on the top and on the right of each graph. The left three columns are biological replicates from 76 cm and the right three from 30 cm soil. Each row corresponds to an incubation time (2 h to 168 h). The BONCAT+ cells gate is displayed as a gray box in each plot, and the percent cells in the BONCAT+ gate is indicated in the top left corner of the box.

**Table S1:** Sample description, pooling strategy and alpha diversity metric of the iTag libraries

| Sample ID | Sample depth | Incubation time | Type | Number of cells pooled | Frequency* per sample before filtering contaminants | Frequency* per sample without contaminant | ESV count | OTU count (97% sim. cluster) |
| --- | --- | --- | --- | --- | --- | --- | --- | --- |
| 30cm_48h_A_BONCAT+ | 30 cm | 48h | BONCAT + | 50000 | 258672 | 257606 | 138 | 97 |
| 30cm_48h_A_BONCAT- | 30 cm | 48h | BONCAT - | 75000 | 140530 | 136389 | 77 | 51 |
| 30cm_48h_B_BONCAT+ | 30 cm | 48h | BONCAT + | 75000 | 236375 | 234877 | 203 | 144 |
| 30cm_48h_B_BONCAT- | 30 cm | 48h | BONCAT - | 75000 | 82431 | 81186 | 171 | 123 |
| 30cm_48h_D_BONCAT+ | 30 cm | 48h | BONCAT + | 75000 | 279152 | 278404 | 178 | 126 |
| 30cm_48h_D_BONCAT- | 30 cm | 48h | BONCAT - | 75000 | 155187 | 89480 | 139 | 105 |
| 76cm_2h_A_BONCAT+ | 76 cm | 2h | BONCAT + | 35000 | 163551 | 163183 | 194 | 131 |
| 76cm_2h_A_BONCAT- | 76 cm | 2h | BONCAT - | 50000 | 122347 | 120373 | 74 | 53 |
| 76cm_2h_B_BONCAT+ | 76 cm | 2h | BONCAT + | 65000 | 240574 | 240285 | 199 | 143 |
| 76cm_2h_B_BONCAT- | 76 cm | 2h | BONCAT - | 75000 | 154938 | 152259 | 55 | 42 |
| 76cm_2h_C_BONCAT+ | 76 cm | 2h | BONCAT + | 73000 | 149499 | 149499 | 268 | 187 |
| 76cm_2h_C_BONCAT- | 76 cm | 2h | BONCAT - | 75000 | 115293 | 115293 | 166 | 109 |
| 76cm_48h_A_BONCAT+ | 76 cm | 48h | BONCAT + | 75000 | 736900 | 736231 | 161 | 129 |
| 76cm_48h_A_BONCAT- | 76 cm | 48h | BONCAT - | 45000 | 146726 | 130866 | 57 | 38 |
| 76cm_48h_B_BONCAT+ | 76 cm | 48h | BONCAT + | 75000 | 257443 | 257311 | 403 | 269 |
| 76cm_48h_B_BONCAT- | 76 cm | 48h | BONCAT - | 35000 | 127528 | 113817 | 126 | 87 |
| 76cm_48h_C_BONCAT+ | 76 cm | 48h | BONCAT + | 50000 | 484680 | 482851 | 613 | 441 |
| 76cm_48h_C_BONCAT- | 76 cm | 48h | BONCAT - | 75000 | 164342 | 160961 | 135 | 99 |
| 76cm_A_filter | 76 cm | - | FILTER | - | 305866 | 305866 | 528 | 381 |
| 76cm_B_filter | 76 cm | - | FILTER | - | 336054 | 336054 | 561 | 421 |
| 76cm_C_filter | 76 cm | - | FILTER | - | 175457 | 175457 | 373 | 281 |
| 30cm_A_filter | 30 cm | - | FILTER | - | 327415 | 327415 | 567 | 406 |
| 30cm_B_filter | 30 cm | - | FILTER | - | 260298 | 260298 | 412 | 296 |
| 30cm_C_filter | 30 cm | - | FILTER | - | 141273 | 141273 | 338 | 246 |
| 76cm_total_soil | 76 cm | - | SOIL | - | 349341 | 349341 | 390 | 324 |
| 30cm_total_soil | 30 cm | - | SOIL | - | 314201 | 314201 | 916 | 620 |

\*Number of sequences after QC and denoising, note that DADA2 removed all singletons

